# Understanding gene regulatory mechanisms based on gene classification

**DOI:** 10.1101/2021.10.31.466641

**Authors:** Hao Tian, Yueying He, Yue Xue, Yi Qin Gao

## Abstract

The CpG dinucleotide and its methylation play vital roles in gene regulation as well as 3D genome organization. Previous studies have divided genes into several categories based on the CpG intensity around transcription starting sites (TSS) and found that housekeeping genes tend to possess high CpG density while tissue-specific genes are generally characterized by low CpG density. In this study, we investigated how the CpG density distribution of a gene affects its transcription and regulation pattern. Based on the CpG density distribution around TSS, the human genes are clearly divided into different categories. Not only sequence properties, these different clusters exhibited distinctly different structural features, regulatory mechanisms, and correlation patterns between expression level and CpG/TpG density. These results emphasized that the usage of epigenetic marks in gene regulation is partially rooted in the sequence property of genes, such as their CpG density distribution.

## Introduction

It is well known that the distribution of CpG dinucleotide is uneven along the genome sequence(Antequera, 2003). For instance, CpGs could accumulate to form the CpG island (CGI)(Antequera, 2003; Deaton and Bird, 2011). The CpG density around the transcription start site (TSS) is generally higher than its surrounding sequence given that the majority of gene promoters are associated with CGI(Deaton and Bird, 2011; Vavouri and Lehner, 2012). Based on the normalized CpG density/GC content of promoter regions, genes have been divided into two or three categories, including HCP (high-CpG promoter), LCP (low-CpG promoter) and ICP (promoters with intermediate CpG contents)(Couldrey et al., 2014; Hartung et al., 2012; Saxonov et al., 2006; Weber et al., 2007; Yang et al., 2014). HCP genes tend to be housekeeping genes whereas LCP genes are more likely to be tissue specific(Saxonov et al., 2006; Yang et al., 2014). These classifications were normally performed based on the average CpG intensity. Since the distributions of CpG can also vary greatly within genes, it is interesting to explore whether the distribution in and around a gene affects its regulatory mechanisms (such as the deposition of epigenetic marks) and function.

The methylation of the cytosine of CpG represents a very important epigenetic mark which appears to be also dependent on the density distribution of CpG along the genome. The methylation level of CGI is usually low, especially in the promoter regions of highly expressed genes(Aoto et al., 2020; Deaton and Bird, 2011; Vavouri and Lehner, 2012). Besides, polycomb repressive complex 2 (PRC2), which is thought to participate in the methylation of lysine on H3, tends to bind to CpG-dense regions under the help of Polycomb-like proteins(Li et al., 2017; van Kruijsbergen et al., 2015). The mutual exclusion between CGI methylation and tri-methylation of H3K27 was observed in both human and mice cells(Bogdanović et al., 2011; Lynch et al., 2012). H3K27me3-marked DNA methylation canyons can form long-range chromatin interactions, which was associated to specific gene repression(Zhang et al., 2020). In general, epigenetic marks, including DNA methylation, active histone marks H3K4me3, H3K36me3 and H3K27ac, as well as repressive histone marks H3K27me3 and H3K9me3, are related to gene activation or repression(Benayoun et al., 2014; Heintzman et al., 2007; Jang et al., 2017; Ninova et al., 2019; Santos-Rosa et al., 2002). Notably, at the same time, many studies also showed the decoupling of epigenetic marks from gene regulation(Borsari et al., 2020; Murray et al., 2019). It is thus intriguing to investigate the possible reasons behind such inconsistency, in particular, whether the DNA sequence plays a role in the usage of different epigenetic regulation.

To address the questions described above, we performed in this paper gene classification based on the CpG density distribution around TSS using Recurrent Neural Network (RNN). The analyses revealed that, in general, human genes can be divided into three categories. Cluster 1 genes possess a high and sharp CpG density peak around TSS. Cluster 2 genes harbor a lower but broader CpG density peak, compared to cluster 1 genes. In contrast, cluster 3 genes are characterized by low CpG densities around TSS. Not only the sequence property, the patterns of nucleosome occupancy and transcription factor (TF) binding, the correlation between gene expression level and CpG/TpG density, the regulatory mechanisms are also distinctly different among the three gene clusters. The promoter regions of cluster 1 genes tend to reside in an open chromatin environment. Cluster 3 genes are more likely to locate in a compact environment and our further analysis revealed that their activation involves the participation of specific transcription factors (TFs), which may facilitate their movement to an euchromatin environment. Besides, genes exhibiting expression variations uncoupled with histone modification variations tend to be in cluster 3 while the expression regulation of cluster 2 genes are most correlated with H3K27me3 among all three classes. In tumorigenesis, the variations of expression and epigenetic marks were also distinctly different for the three gene clusters. Together, our results emphasized the importance of taking the genetic sequence properties into account for understanding the gene regulatory mechanisms.

## Results

### Gene classification based on CpG distribution

Many studies have revealed the distinctly different CpG distributions around TSS between housekeeping genes and tissue-specific genes(Roider et al., 2009; Saxonov et al., 2006; Yang et al., 2014). Here, we firstly utilized the Recurrent Neural Network (see Methods) to distinguish most of the housekeeping genes from tissue-specific genes. Briefly, we downloaded the human promoter CAGE data from FANTOM5 project (https://fantom.gsc.riken.jp/5/datafiles/latest/extra/CAGE_peaks/) and regarded the midpoint of CAGE peaks labeled with “p1@” as gene TSS. For each gene, the corresponding CpG density distribution was calculated based on the DNA sequences of 16 kb (8 kb upstream to 8 kb downstream of TSS) using the 40-bp non-overlapping window, thus generating a 1 × 400 vector. The network was first trained (see Methods) using the CpG density distributions of a proportion of housekeeping genes (downloaded from UCSC database) and tissue-specific genes(Tian et al., 2020). The AUC value of the testing set is 0.947.

Next, we applied this well-trained model to the entire gene set, each gene was therefore conferred by a “CG likelihood”, the value of which indicates the confidence that one gene could be regarded as a tissue-specific gene. Similar to previous studies(Weber et al., 2007), the distribution of CG likelihood roughly consisted of three parts (Figure 1A), indicating that the human genes could be divided into three clusters and we performed such a classification by means of Gaussian Mixture Model. Genes in clusters 1 (gene number is 12361) and 3 (gene number is 8396) possess the lowest and highest CG likelihood, and are characterized by highest (and sharp) and lowest CpG density distribution around TSS, respectively, whereas cluster 2 genes (gene number is 3752) have broad CpG peaks around TSS featured by moderate intensity (Figure 1B). Accordingly, almost all (3123 out of 3447) housekeeping genes are in cluster 1, and in contrast, the majority of (1292 out of 1821) tissue-specific genes belong to cluster 3. Genes in clusters 1 and 3 harbor the lowest and highest tissue specificities (higher value of this parameter indicates the corresponding genes are specifically and highly expressed in fewer tissues(Tian et al., 2020), see Methods), respectively (Figure 1C), accordant with previous studies revealing the negative correlation between TSS CpG density and tissue specificity score(Yang et al., 2014). Notably, although tissue-specific genes tend to possess low CpG densities around TSS, most tissue-specific transcription factors (ts-TFs) are relatively CpG-rich, given that 108 out of 171 ts-TFs are in cluster 2. We also analyzed the distribution of tissue-specific genes pertinent to certain tissues (e.g., liver-specific genes) among three clusters. Interestingly, brain-specific genes tend to reside in clusters 1 and 2, while liver, spleen, and whole blood-specific genes are more likely to belong to cluster 3 (Figure S1A).

**Figure 1.**
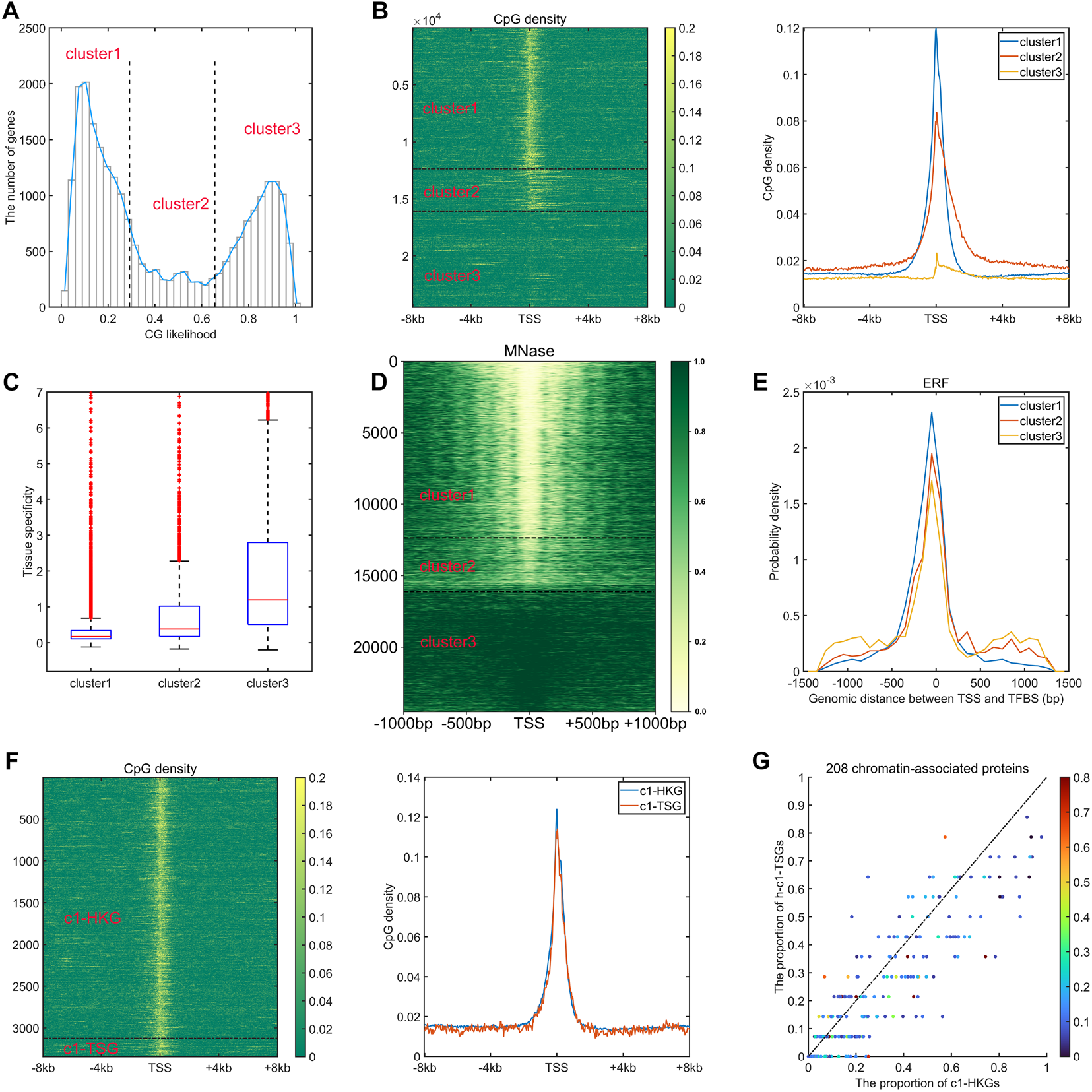
Gene classification based on CpG distribution. **(A)** The distribution of CG likelihood of human genes. **(B)** The CpG density distribution of genes belonging to different clusters (left, heatmap; right, average behaviors). **(C)** The tissue specificity of genes of three clusters. Based on the definition of tissue specificity (see Methods), for each gene, its maximum tissue specificity among 37 tissues (testis was not considered since it contains too many tissue-specific genes) was extracted and drawn here. **(D)** The nucleosome occupancy patterns (measured by MNase-seq) of genes of three clusters in GM12878. **(E)** The distribution of genomic distance between TF binding sites and gene TSS. Positive and negative values indicate that TFs bind to the regions downstream and upstream of TSS, respectively. **(F)** The CpG density distribution of c1-HKG and c1-TSG (left, heatmap; right, average behaviors). **(G)** The proportion of c1-HKGs/h-c1-TSGs that bind to one certain TF. Each data point represents one certain TF, the corresponding color represents its tissue specificity. P-value = 4.12 × 10^−11^ by t-test.

Akin to CpG density distribution, nucleosome occupancy patterns are also found to be significantly different among the three clusters. In general, as for cluster 1 genes, the promoter regions of which reside in an open environment and the arrangement of nucleosome downstream of TSS is regular (Figure 1D). Such a feature can also be observed for cluster 2 genes, but to a much lesser extent (Figure 1D). By contrast, cluster 3 genes are generally located in a compact and nucleosome-occupied environment, especially for promoter regions (Figure 1D). A recent study has uncovered the binding sites (ChIP-seq peaks) of 208 chromatin-associated proteins (CAPs), including 171 TFs and 37 transcriptional cofactors and chromatin regulator proteins, in HepG2 cell(Partridge et al., 2020). We found that the binding sites of the majority of CAPs, e.g., ERF, ELF3, CHD2, MAZ, tend to reside in a specific region for cluster 1 genes, while the binding patterns become more “dispersed” for clusters 2 and 3 genes (Figure 1E and S1B).

The housekeeping genes and tissue-specific genes in cluster 1 (named c1-HKGs and c1-TSGs, respectively, although the number of the latter is relatively low: 3123 vs 221) possess very similar CpG distribution patterns (Figure 1F) and they both locate in an open environment (Figure 1D). To understand the factors contributing to their different transcriptional activities, we examined their CAP binding patterns. Since in contrast to c1-HKGs, many c1-TSGs are not expressed in HepG2, we chose as examples the highly expressed c1-TSGs (named h-c1-TSGs), the expression level of which exceeds the 75^th^ percentile of all genes’ expression level and is comparable to c1-HKGs (p-value > 0.05). One can see from Figure 1G that CAPs are inclined to bind to the promoter regions of c1-HKGs, hinting that the sequence motif of c1-HKGs promoter regions is more likely to recruit TFs. We also compared the CAP binding for highly-expressed c1-TSGs and c3-TSGs and found that CAPs with higher tissue-specificities have a higher tendency to bind to c3-TSGs (Figure S1C), which means that these CAPs are prone to regulate tissue-specific genes with low but not high promoter CpG densities. Aside from sequence features, we also examined the 3D chromatin structure properties and found that the insulation score (one parameter used to assess the possibility that the locus locates in TAD boundary, see Methods) of c1-HKGs is significantly higher than c1-TSGs (Figure S1D). This result shows that compared to c1-TSGs, c1-HKGs are more likely to reside at TAD boundaries, accordant with previous findings that HKGs are enriched near TAD boundaries(Dixon et al., 2012). Together, these results revealed that although the CpG distributions around the TSS are similar, c1-HKGs and c1-TSGs display distinctly different sequence and structure traits. The sequence properties for one gene to be a HKG include not only the high CpG density within the promoter region, which permits the open chromatin structure (Figure 1D), but also the specific sequence motif which can effectively recruit different kinds of TFs for transcription.

We note here that there exist many studies(Couldrey et al., 2014; Hartung et al., 2012; Saxonov et al., 2006; Weber et al., 2007; Yang et al., 2014) dividing the gene promoters into two or three categories, for example HCPs (high-CpG promoters), LCP (low-CpG promoters) and ICP (promoters with intermediate CpG content). However, the criterion behind these classifications was based on the (normalized) promoter CpG/G+C content, and did not consider explicitly the CpG density distribution upstream and downstream of TSS. Here, as an example, we compared the CpG distribution as well as the CpG intensity of housekeeping genes and tissue-specific genes around TSS within each cluster identified by RNN and Weber et al(Weber et al., 2007). Minor difference existed within clusters identified by RNN (Figure S1E and S1F), indicating that our classification does provide opportunities to investigate how sequence property variation affects gene regulatory mechanisms. In fact, genes within each cluster (identified in this work) exhibit particular regulatory mechanisms that are significantly different from each other, which will be discussed below.

### The relation between sequence property and gene expression

Earlier studies revealed a positive correlation between CpG density around TSS and gene expression in vertebrate(Cheng et al., 2012; Yang et al., 2014). This result is not surprising since housekeeping genes are characterized by high CpG density in promoter regions and are normally highly expressed. Here, we asked whether such a positive correlation between expression and CpG density still holds in each individual cluster. We calculated the Spearman correlation coefficient between gene expression level and CpG density of each non-overlapping 40-bp window among genes belonging to the same cluster (see Methods) and found that CpG densities around TSS are in general positively correlated with gene expression, regardless of the cluster under study (Figure 2A, S2 and S3). Furthermore, we noticed that this correlation is higher for clusters 2 and 3 genes than for cluster 1 genes in most samples we examined (including early embryonic cells, somatic cells, tumor cells and its corresponding paracancerous cells, Figure 2A, S2 and S3). In a few samples (e.g., early embryonic cells, hES, LIHC, BLCA), we did not observe the prominent correlation peak near the TSS (Figure 2A and S3) for cluster 1 genes. The correlation level downstream of TSS (TSS - ∼+4kb) decays much slower for cluster 3 and is thus more positive than clusters 1 and 2 genes (Figure 2A and S2). These results thus show that aside from sequence properties, the correlation patterns derived from different gene clusters are also significantly different (therefore the correlation pattern may be influenced intrinsically by sequence properties).

**Figure 2.**
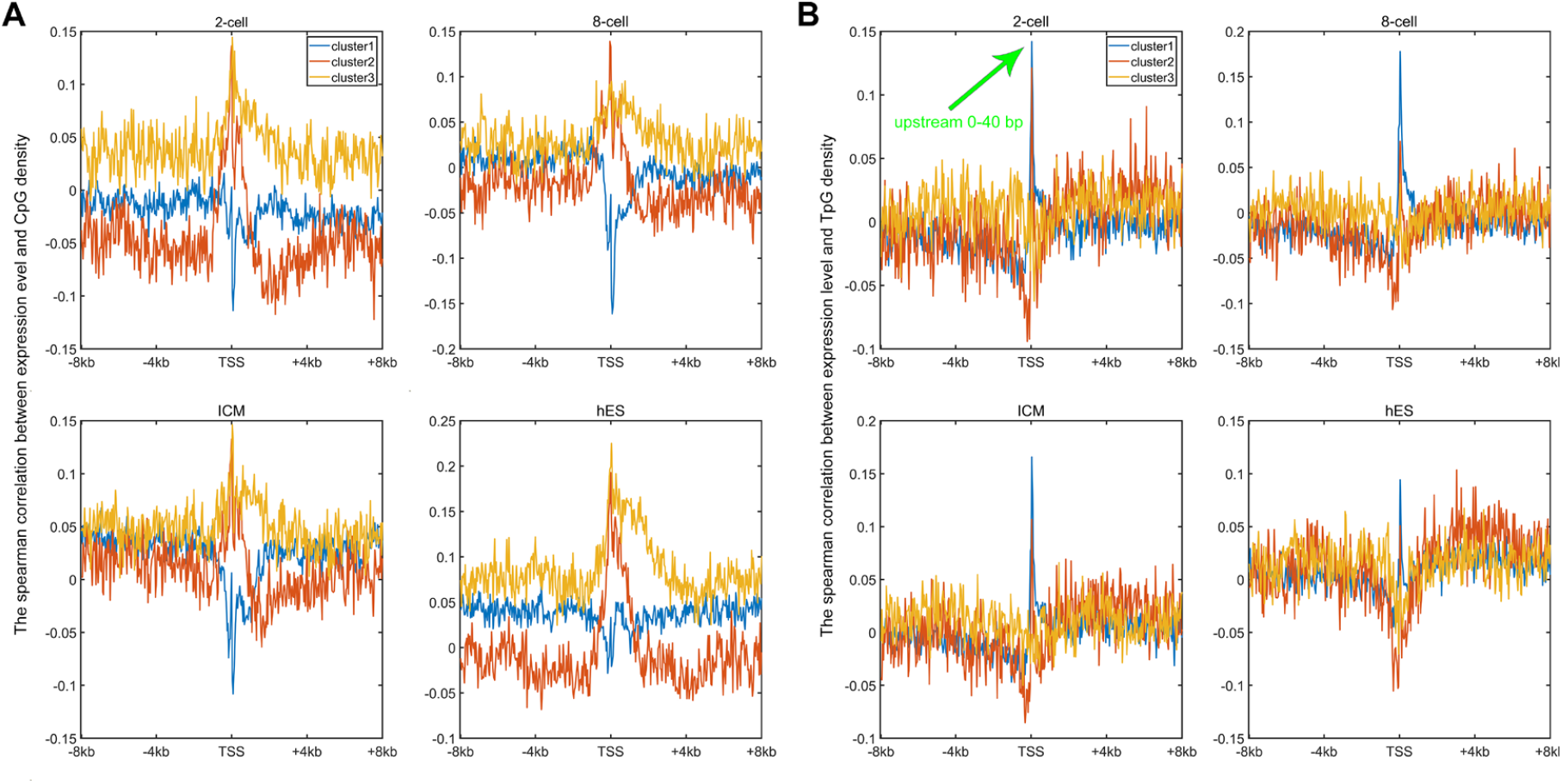
The relation between sequence property and gene expression. **(A)** The Spearman correlation between gene expression and CpG density of each non-overlapping 40 bp window (see Methods). **(B)** The Spearman correlation between gene expression and TpG density of each non-overlapping 40 bp window.

An earlier study revealed that the expression level was negatively correlated with TpG density near TSS of pig genes (Schachtschneider et al., 2015). We found here that in different human cells, these three clusters exhibit an overall negative correlation between expression and the TpG density near TSS, but this correlation level for clusters 2 and 3 genes is generally more negative than cluster 1 genes (Figure 2B, S4 and S5). Intriguingly, as for clusters 1 and 2, the correlation between expression and TpG density 0∼40bp downstream of TSS is positive and thus noticeably different from its surrounding sequences (Figure 2B, S4 and S5). In the following, we show that this region also distinguishes itself by unique histone modification marks.

### Distinct regulatory mechanisms among three gene clusters

Since the three gene clusters have distinct sequence features and epigenetic mark usage is expected to be affected by DNA sequence, we next investigated whether the corresponding epigenetic mechanisms also differ for the three classes of genes. We analyzed the RNA-seq, histone modification and DNA methylation data for liver, lung, ovary and sigmoid colon. Similar results were obtained for each set of data. Taking liver as an example, only a small proportion of cluster 1 genes are marked with repressive histone modifications (H3K27me3 and H3K9me3, Figure 3A and S6A) and as expected, these genes possess relatively low expression levels within cluster 1 (Figure 3A and S6A). Most cluster 1 genes are decorated with active histone marks (H3K4me3 and H3K36me3), consistent with their higher expression levels (Figure 3B and S6B). Intriguingly, one feature for repressed genes of cluster 2 is that their high H3K27me3 signals tend to be dispersed along the genome sequence upstream and downstream of TSS (Figure 3A). In contrast, the H3K27me3 distributions of cluster 1 repressed genes are limited close the TSS (Figure 3A). For cluster 3 genes, although a higher proportion of genes are repressed, compared to clusters 1 and 2, a dispersed and much weaker pattern of H3K27me3 is observed for repressed genes (Figure 3A), indicating that the repression of cluster 3 genes may be largely independent of the deposition of H3K27me3. Such results are accordant with previous studies revealing that PRC2, playing a vital role in trimethylation of H3 on lysine K27, tends to bind to CG-rich regions(Li et al., 2017; Mendenhall et al., 2010; van Kruijsbergen et al., 2015).

**Figure 3.**
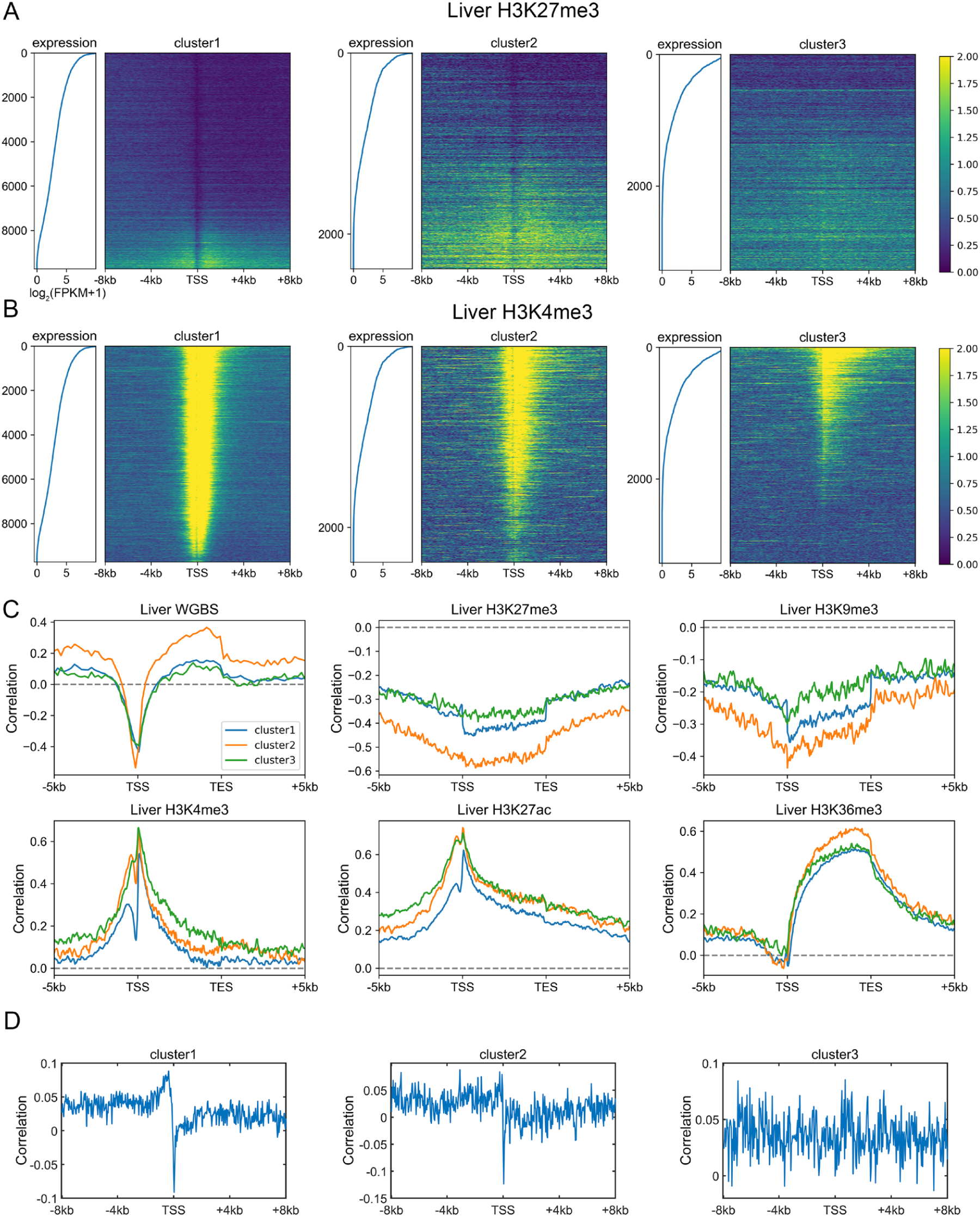
Distinct regulatory mechanisms among three gene clusters. **(A)**-**(B)** The distribution of H3K27me3 **(A)** and H3K4me3 **(B)** among three gene clusters in liver. Each heatmap was ranked based on gene expression level. **(C)-(D)** The Spearman correlation between gene expression level and epigenetic marks **(C)**, and between TpG density and H3K27me3 **(D)** (see Methods).

To quantify the expression dependences on epigenetic modifications among genes with different DNA sequences, akin to the correlation between CpG density and gene expression level introduced above, for each cluster, we calculated the spearman correlation between expression levels and epigenetic marks around the TSS for different genes (Figure 3C and S7). The correlation between the expression level and DNA methylation is negative near TSS, and is positive both up- and down-stream of TSS for all three types of genes (Figure 3C). Notably, among all three gene clusters, the expression level of cluster 2 genes is most strongly correlated to DNA methylation. As for repressive histone modifications (H3K27me3 and H3K9me3), the gene expression is negatively correlated with modification level, especially for cluster 2 genes (Figure 3C). Interestingly, we found that the transcription factor EZH2(Partridge et al., 2020), which is thought to be involved in the methylation of H3K27, tends to bind to the promoter regions of cluster 2 genes (p-value= 6.67 × 10^−20^ for cluster 1 and 2, p-value = 3.42 × 10^−76^ for cluster 2 and 3, Fisher’s exact test), indicating that the wide usage of repressive histone mark for the downregulation of cluster 2 genes. For active histone modifications, as expected, the expression is positively correlated with H3K4me3 and H3K27ac signals around TSS, and with H3K36me3 signal in gene body (Figure 3C). These correlation levels are generally lower for cluster 1 genes than for genes of the other two clusters (Figure 3C).

The above results revealed that different gene clusters exhibit distinct correlation patterns between expression level and epigenetic marks, hinting the important role of DNA sequence feature in the aspect of epigenetics. In fact, as we introduced above, the H3K27me3 distribution of repressed genes belonging to different clusters does correlate with the corresponding CpG density distribution. In fact, cluster 3 genes tend not to be occupied by H3K27me3 for repression, resulting in the low correlation between H3K27me3 and expression level. In contrast, cluster 2 genes tend to recruit proteins responsible for trimethylation of H3K27 (such as polycomb group proteins) for repression due to their sequence characteristics (likely, the high and broad CpG density peak), leading to the higher correlation between expression and H3K27me3 signal. A recent study(Borsari et al., 2020) investigated the variation of expression and nine histone modification signals of human coding genes along the transdifferentiation process, from pre-B cell to macrophage, and identified a gene cluster: genes within which are not marked by most kinds of histone modifications throughout the differentiation process but exhibit expression variation. Intriguingly, the majority (70%) of these genes are found here to belong to cluster 3, indicating that the expression level of genes characterized by low-CpG density are more likely uncouple from histone marks. To gain information on the regulatory mechanism of cluster 3 genes, we calculated the correlation between compartment index (gene with higher value indicates it locates in a more compartment A environment, see Methods) and gene expression level among different tissues and found that this correlation for cluster 3 genes appears to be higher than other two types of genes (Figure S6C). Such a result indicates that chromatin structure organization plays a more important role in cluster 3 gene regulation: these genes are intrinsically prone to silence and their activation are likely facilitated by a movement to a more compartment A-like environment, possibly under the help of specific transcription factors(Boija et al., 2018; Hnisz et al., 2017; Hnisz and Young, 2017; Kim and Shendure, 2019; Stadhouders et al., 2019; Tian et al., 2020). As mentioned above, cluster 1 and 2 genes show a sharp-positive peak in the correlation between expression level and TpG density 0∼40 bp downstream of TSS (Figure 2B). Consistently, we found a salient negative correlation between TpG density 0∼40 bp downstream of TSS and repressive histone modifications (H3K27me3, Figure 3D).

### Expression and epigenetic changes in carcinogenesis

Given that gene dysregulation is a hallmark of cancer(Hanahan and Weinberg, 2011), we next investigated whether genes belonging to different clusters show different expression variations in carcinogenesis. In general, during tumorigenesis, housekeeping-like cluster 1 genes tend to be upregulated, while cluster 3 genes show an opposite tendency (Figure 4A). This result is consistent with our previous finding that genes with high CpG densities tend to be upregulated in cancer cells(Xue et al., 2020), indicating that the expression change in carcinogenesis is partially coupled with the CpG density. Nevertheless, Figure 4A shows that although cluster 1 genes are more likely to be up-regulated, the changes tend to be small. By comparison, cluster 3 genes display a broad distribution of expression fold change (Figure 4A), suggesting that some genes are dramatically activated/repressed in tumorigenesis. In addition, consistent with our previous findings(Xue et al., 2020), genes specifically and highly expressed in paracancerous tissue (mainly in cluster 3) are mostly down-regulated in cancer cells (Figure S8A) whereas complementary genes (that is, genes specifically expressed in other tissues other than paracancerous) are mostly up-regulated (Figure S8B).

**Figure 4.**
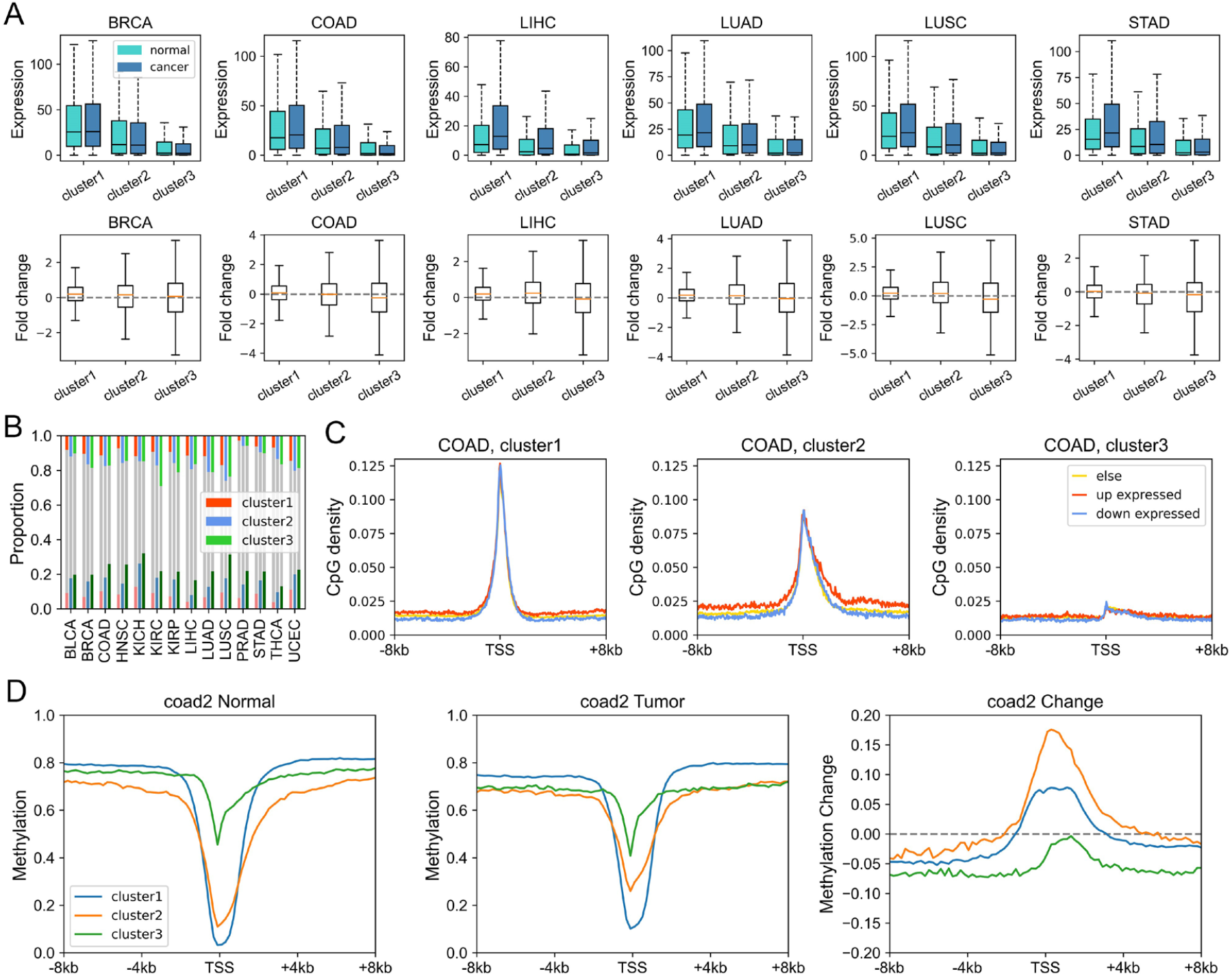
Expression and epigenetic changes in carcinogenesis. **(A)** The expression level (TPM) of genes of different clusters in cancer and normal samples (upper) and the log_2_(expression fold change) in carcinogenesis calculated by Deseq2 (down). **(B)** The proportion of DE genes (in carcinogenesis) in three clusters. Upper: the proportion of up-expressed genes, down: the proportion of down-expressed genes. **(C)** The CpG density distribution of up-expressed, down-expressed and other genes. COAD (colon cancer) was used here for illustration. See Figure S9 for more instances. **(D)** The DNA methylation level of genes of different clusters in normal (left) and tumor (middle) cells, and the methylation changes during carcinogenesis for three gene clusters (right). COAD was used for illustration. See Figure S11 for more instances.

Next, we investigated the DNA sequence characteristics, functions and epigenetic modifications of genes, the expression variations of which are large in carcinogenesis (hereafter referred to as DE genes, which are identified by Deseq2(Love et al., 2014), see Methods). The proportions of DE genes in clusters 1 and 3 are lowest and highest, respectively, in all cancer types (Figure 4B), consistent with the above results that expressions of cluster 1 genes change the least, and those of class 3 genes, the most, in carcinogenesis. The CpG density up- and down-stream of TSS for up-regulated genes are slightly higher than other genes almost for all types of cancers, especially for cluster 2 genes (Figure 4C and S9). GO function analyses show that up-regulated genes in cluster 1 are closely related to cell cycle, including nuclear division, DNA replication and chromosome segregation, contributing to the uncontrolled growth of tumor (Figure S10). Down-expressed genes in clusters 1 and 3 are often related to the muscle system process (Figure S10). The DE genes in cluster 2 are found to be associated with development, including embryonic organ morphogenesis, cell fate commitment (Figure S10), which were suggested to relate to the dedifferentiation and invasion of cancer cells(Ma et al., 2010).

On the other hand, we found that genes enriched in H3K27me3 in paracancerous (normal) tissue, which is a standing-out property for developmental genes(Schuettengruber et al., 2017), tend to become dysregulated in cancer cells, especially for cluster 2 genes. To be specific, for colon cancer, compared with stably expressed genes (in carcinogenesis), up- and down-expressed genes have significantly higher H3K27me3 levels in their corresponding normal tissues (Figure S11A). At the same time, the TSS of these genes are also likely to be hypermethylated in carcinogenesis (Figure S11A). Traditionally, DNA hypermethylation around TSS is associated with the gene repression, this hypermethylation-associated gene activation in cancer is possibly a result of their repression by a broad H3K27me3 signal around TSS in normal cells (Figure S11A). Not only the TSS, but also the upstream and downstream of TSS of these up-expressed genes are hypermethylated in cancer cells (Figure S11A), with the latter being favorable for gene expression (Luo et al., 2018; Schachtschneider et al., 2015). In addition, the TSS of cluster 2 genes undergo the most significant hypermethylation (Figure 4D, S11B-D) among the three types of genes, consistent with them being the most enriched in H3K27me3 in normal cells. Finally, the up- and down-stream of TSS of cluster 3 genes tend to be more hypomethylated than other genes (Figure 4D, S11B-D), which is likely because that cluster 3 genes are CpG-poor and mainly located in the repressive compartment B, in which hypomethylation is more likely to be found.

## Discussion

To investigate how much CpG density distribution of a gene affects its expression and regulation pattern, we performed in this study the gene classification using recurrent neural network. We then analyzed the biological functions and the regulatory mechanisms of genes with different sequence properties. Compared to previous methods, which classified genes based on CpG/C+G intensity of promoter regions(Saxonov et al., 2006; Weber et al., 2007), our classification considered the CpG distribution in a large region around the TSS and thereby included more comprehensive sequence features. The model yielded three gene clusters, each with considerably different sequence features. We found that three gene clusters are distinctly different in terms of expression, regulatory mechanisms, chromatin structural features, and TF binding patterns. For instance, cluster 1 genes tend to reside intrinsically in an open chromatin environment and their activation relies weakly on specific genome reorganization. In contrast, the promoter regions of cluster 3 genes have low CpG densities and accessibility. Tissue-specific transcription factors (such as pioneer factors(Iwafuchi-Doi and Zaret, 2014), as discussed below) may play a vital role in the regulation of cluster 3 genes given that these factors could overcome the nucleosome barriers and tether the cluster 3 genes to a more active environment (e.g., through a phase separation mechanism)(Hnisz et al., 2017). Notably, our gene classification clearly identified a gene cluster, the promoter regions of genes within which are characterized by a high and broad CpG peak (cluster 2). The regulation of cluster 2 genes was found to be most strongly correlated with epigenetic modifications, especially for H3K27me3, among all three types of genes. Given that the association between epigenetic mark and gene expression is still controversial and different gene clusters exhibit different degrees of dependence on epigenetic marks, we speculate that the confusion may, at least in part, result from the diversity of CpG distribution around promoter regions of different genes.

Intriguingly, we found that for a small number of tissue-specific genes (c1-TSGs), the CpG density distribution around TSS is almost the same as that for housekeeping genes. Their difference appears to exist in TF-binding: c1-TSGs are significantly depleted of TF binding sites compared to c1-HKGs, which may prevent the former from being broadly expressed. We also noticed that unlike (general) tissue-specific genes, which are typically characterized by extremely low CpG density around TSS, genes encoding tissue-specific TFs generally possess high CpG densities (e.g., *FOXA1*). Such a sequence property may render a relative open environment for them to be easily accessed, as can be seen from MNase data, and to be expressed upstream of their target tissue-specific genes. A possible regulatory cascade therefore appears in which following the establishment of cell identity, housekeeping genes and genes encoding housekeeping TFs are activated early, following which genes encoding tissue-specific TFs become activated, and, finally, the tissue-specific TFs access the promoter regions of tissue-specific genes to activate them (Figure S1C). Interestingly, the CpG density of regions recruiting tissue-specific TFs is indeed significantly lower than that of the housekeeping TFs (Figure S12).

The tendency of gene expression changes during carcinogenesis also shows sequence biases as various types of cancers exhibit similar changes, including the preferred expression for high-CpG genes. Besides, the functions of dysregulated genes within one specific gene cluster are also similar for different types of cancers. Together with different modes of DNA methylation change for the three cluster genes, the similar epigenetic and expression changes among different types of cancer indicate that the mechanisms of cancer development are at least partially dictated by the sequence properties of genes, therefore pointing to the importance of including sequence properties in deciphering the cancer epigenomes.

## Materials and Methods

### RNN for gene classification

At first, all housekeeping genes (gene number is 1679) and tissue-specific genes (gene number is 1321) are labeled with 0 and 1, respectively, then 800 housekeeping genes and 800 tissue-specific genes were chosen as training set and the remaining are regarded as testing set. The RNN utilized in this study is based on bidirectional gate recurrent unit, with 1 layer and 16 hidden states, followed by a fully connected network with two hidden layers (the number of nodes is 16 and 2, respectively). To make full use of the unlabeled genes, the loss function contained two parts: supervised and unsupervised term. The supervised term measured the binary cross entropy between the target *y* and output *ŷ*, while the unsupervised term measured the clustering consistency between unlabeled original data and augmented data. The optimizer we used is Adam.

We then applied the well-trained network (which could distinguish housekeeping gene from tissue-specific genes based on CpG density distribution) to the entire human gene set and “CG likelihood” for each gene was obtained.

### The definition of tissue specificity

We downloaded the normalized gene expression data of 38 human tissues from https://zenodo.org/record/838734, and the tissue specificity of gene *i* in tissue *t*, 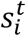, is calculated as(Tian et al., 2020)

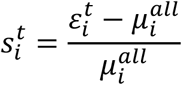

where 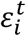 and 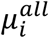 are the mean expression levels of gene *i* in tissue *t* and all tissues, respectively.

### The calculation of insulation score

For each 40-kb bin (the resolution corresponds the resolution of Hi-C matrices in this study), its insulation score (Bintu et al., 2018) *IS* is calculated as

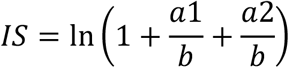

where *a*1 and *a*2 are the average contact probabilities of its two flanking regions *A*1 and *A*2, respectively, and *b* is the average contact probability of the cross region *B* (the element of *B* represents the spatial interaction between *A*1 and *A*2). The window size we used here is 480 kb.

### The calculation of Spearman correlation coefficient between expression level and CpG/TpG density

For each gene cluster, the size of corresponding CpG density matrix is *n* × 400, where *n* represents the gene number in this cluster and 400 is the window number. Accordingly, the size of expression vector in one specific cell is *n* × 1. The Spearman correlation coefficient is then calculated between expression vector and each column of CpG density matrix, yielding a 1 × 400 correlation coefficient vector. The correlation between TpG density and histone modifications, as well as between expression level and histone modifications are calculated in the same way.

**The calculation of compartment index**

Based on the Hi-C contact matrix, the whole genome could be divided into two compartments, A and B(Lieberman-Aiden et al., 2009), with the former being more open and active while the latter mainly corresponding to heterochromatin. The compartment index (*CI*) of bin *i* is then calculated as

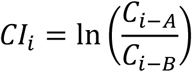

where *C*_*i*−*A*_ and *C*_*i*−*B*_ are the average normalized contact probabilities between bin *i* and compartment A bins, and between bin *i* and compartment B bins, respectively. A higher value of *CI* thus indicates a more open environment.

### The identification of DE genes

During carcinogenesis, genes with log_2_(expression fold change) > 1 and p-value < 0.05 are defined as up-expressed genes, and genes with log_2_(expression fold change) < -1 and p-value < 0.05 are regarded as down-expressed genes. Such results are obtained using Deseq2(Love et al., 2014).

### Gene function analysis

The clusterProfiler package (Yu et al., 2012)was used in this study for gene function analysis.

## Supporting information

Data sources

## Supplementary Materials

Additional file1: supplemental materials. This file contains 12 supplemental figures. Additional file2: data sources. This file summarized the data we used in this study and its sources.

## Author Contributions

Conceptualization, Y.Q.G., H.T., Y.Y.H. and Y.X.; Methodology, Y.Q.G., Y.Y.H., H.T. and Y.X.; Formal Analysis, H.T., Y.Y.H. and Y.X.; Writing – Original Draft, H.T., Y.X. and Y.Y.H.; Writing – Review & Editing, Y.Q.G. and H.T.; Visualization, H.T., Y.X. and Y.Y.H.; Supervision, Y.Q.G.; Funding Acquisition, Y.Q.G.

## Data Availability

The data we analyzed in this work are publicly available and summarized in Additional file 2.

## Acknowledgements

We would like to thank Dr. Zhicheng Cai for insightful discussions.

