## Supplemental figures for "Understanding gene regulatory mechanisms based on gene classification"

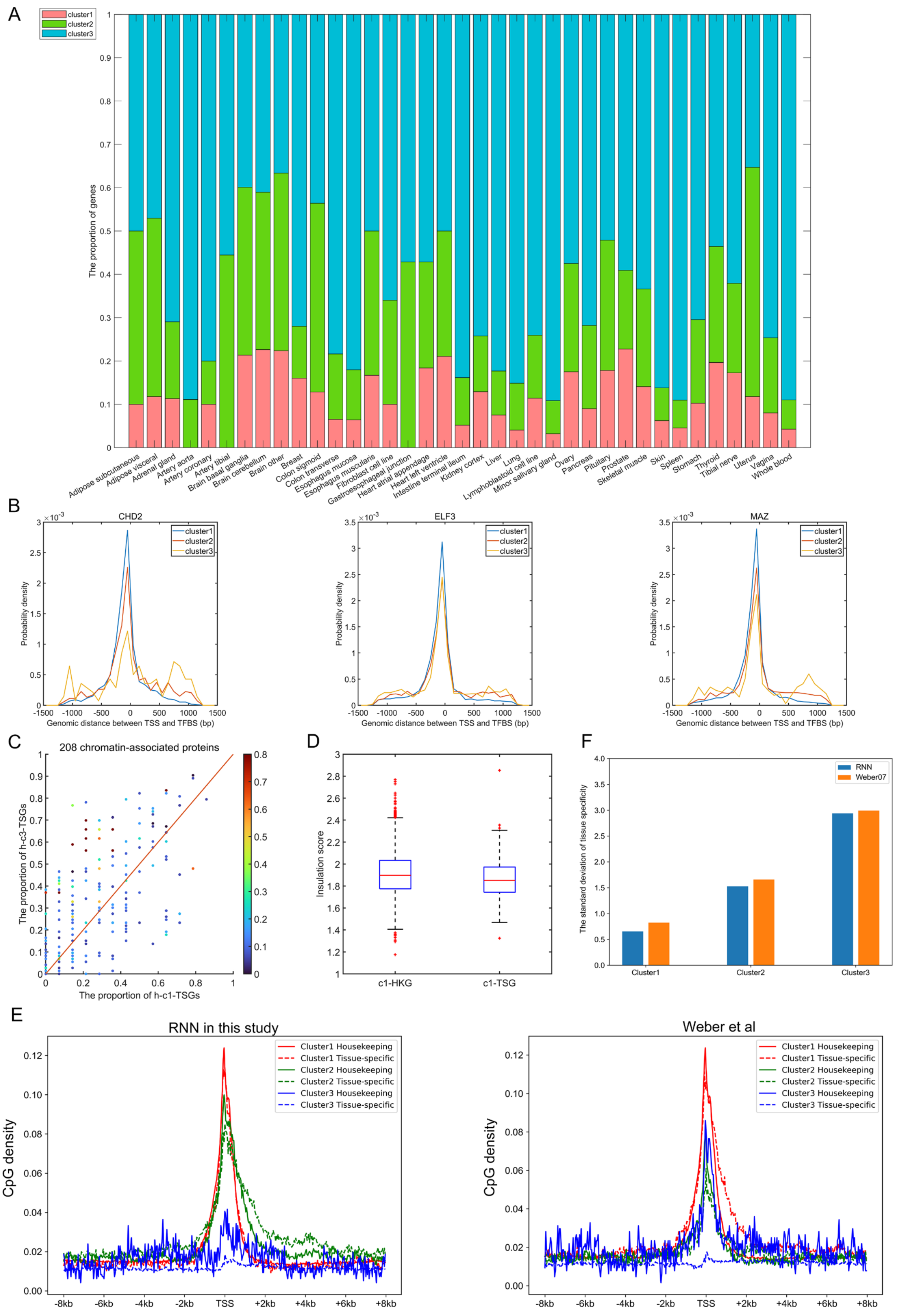


**Figure S1. Gene classification based on CpG distribution. (A)** The distribution of tissue-specific genes of one certain tissue among three clusters. **(B)** The distribution of genomic distance between TF binding sites and gene TSS. Positive and negative values indicate that TFs bind to the regions downstream and upstream of TSS, respectively. **(C)** The proportion of h-c1-TSGs/h-c3-TSGs that bind to one certain TF. Each data point represents one certain TF, the corresponding color represents its tissue specificity. **(D)** The comparison of insulation score between c1-HKGs and c1-TSGs in liver. Window size we used here is 480 kb (see Methods). P-value = $9.4054 \times 10^{-5}$ by Welch's unequal variance test. **(E)** The CpG density distribution of housekeeping genes and tissue-specific genes within each cluster identified by RNN and Weber et al. **(F)** The standard deviation of tissue specificity of each cluster identified by RNN and Weber et al.

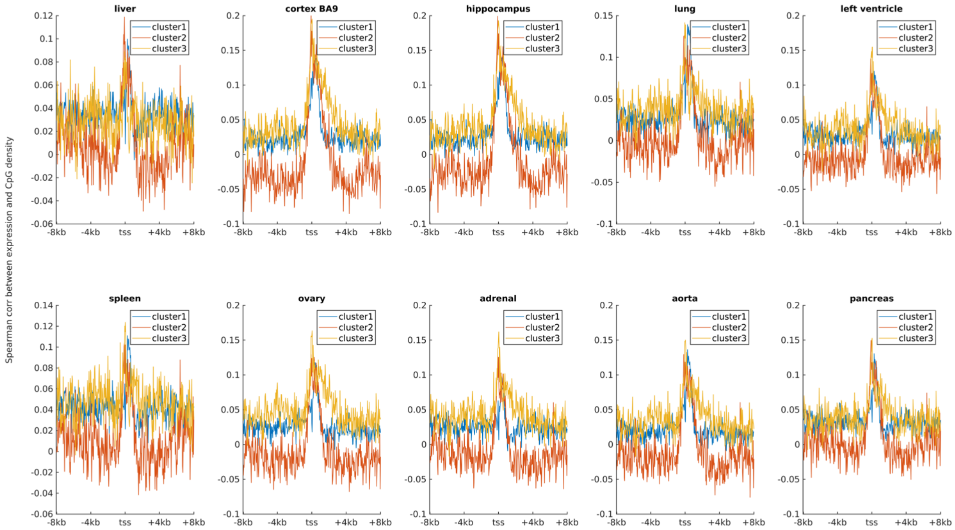


**Figure S2. The Spearman correlation between gene expression level and CpG density in ten human tissues.**

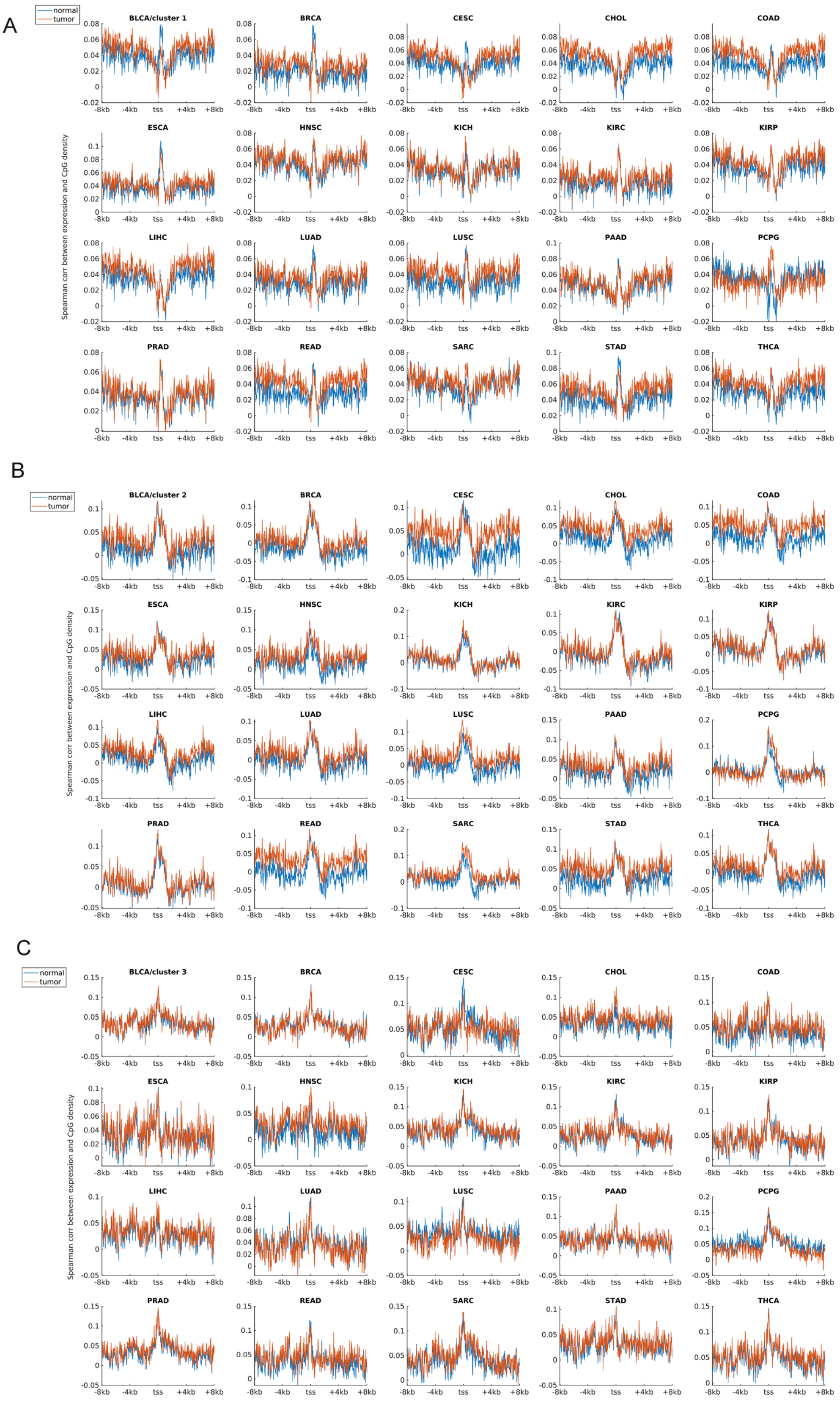


**Figure S3. The Spearman correlation between gene expression level and CpG**

**density in tumor cells and corresponding paracancerous (normal) cells. (A)**, **(B)** and **(C)** represent the correlation in cluster 1, 2 and 3, respectively.

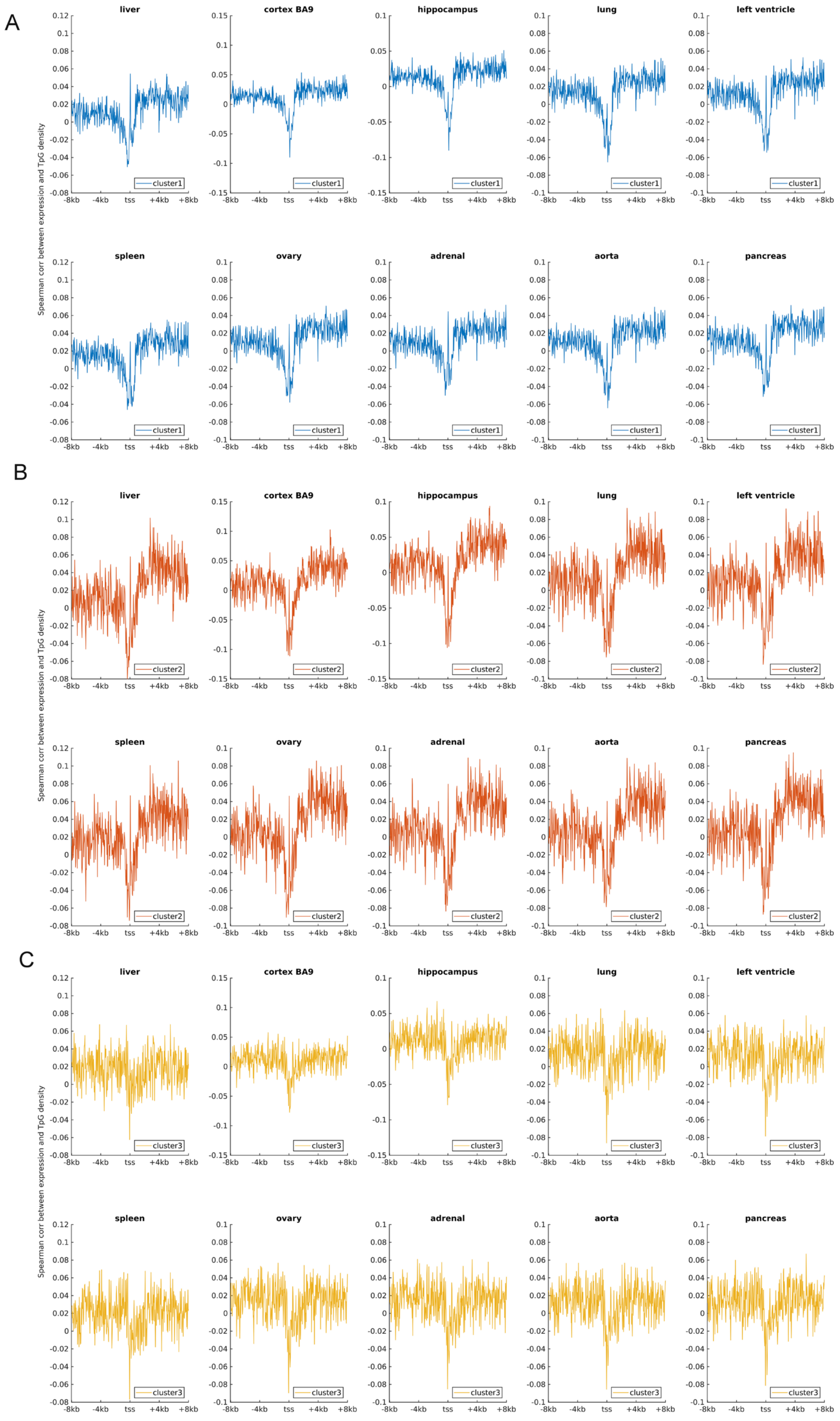


**Figure S4. The Spearman correlation between gene expression level and TpG density in ten human tissues. (A), (B)** and **(C)** represent the correlation in cluster 1, 2 and 3, respectively.

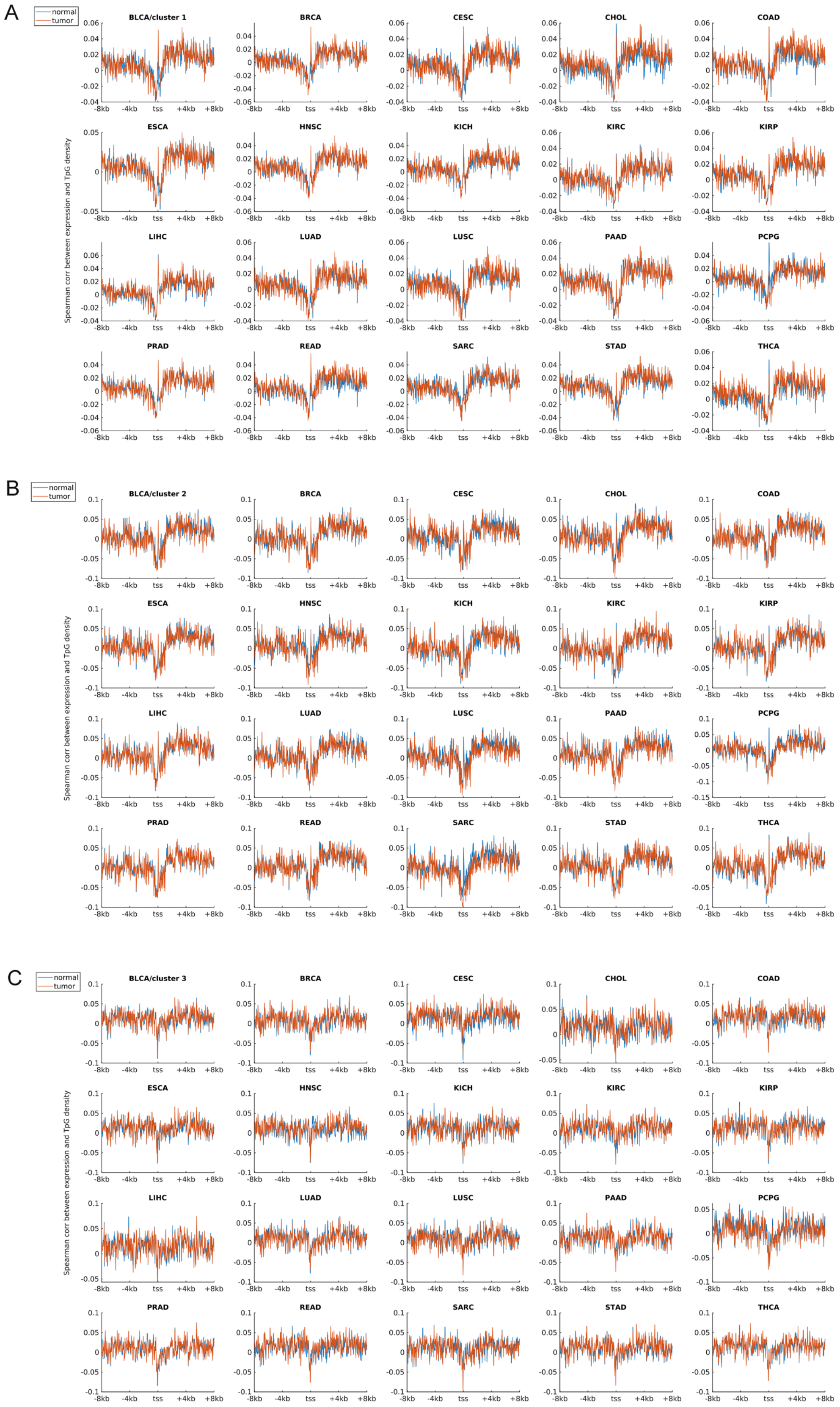


**Figure S5. The Spearman correlation between gene expression level and TpG density in tumor cells and corresponding paracancerous cells. (A), (B)** and **(C)**

represent the correlation in cluster 1, 2 and 3, respectively.

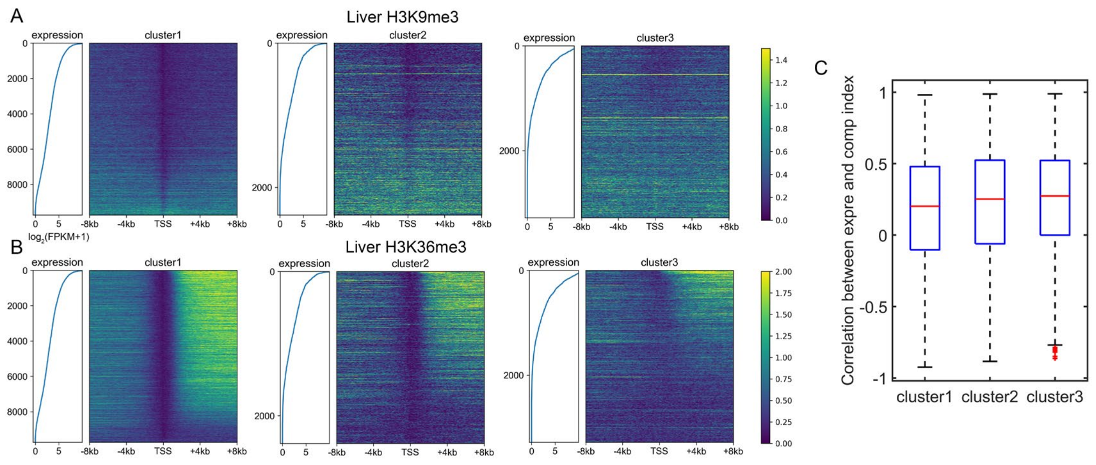


**Figure S6. Distinct regulatory mechanisms among three gene clusters. (A)-(B)** The distribution of H3K9me3 **(A)** and H3K36me3 **(B)** among three gene clusters in liver. Each heatmap was ranked based on gene expression level. **(C)** The Pearson correlation coefficient between compartment index and gene expression level among different tissues. P-value = $1.16 \times 10^{-6}$ for clusters 1 and 2, p-value = $0.0067$ for clusters 2 and 3, Welch's unequal variance test.

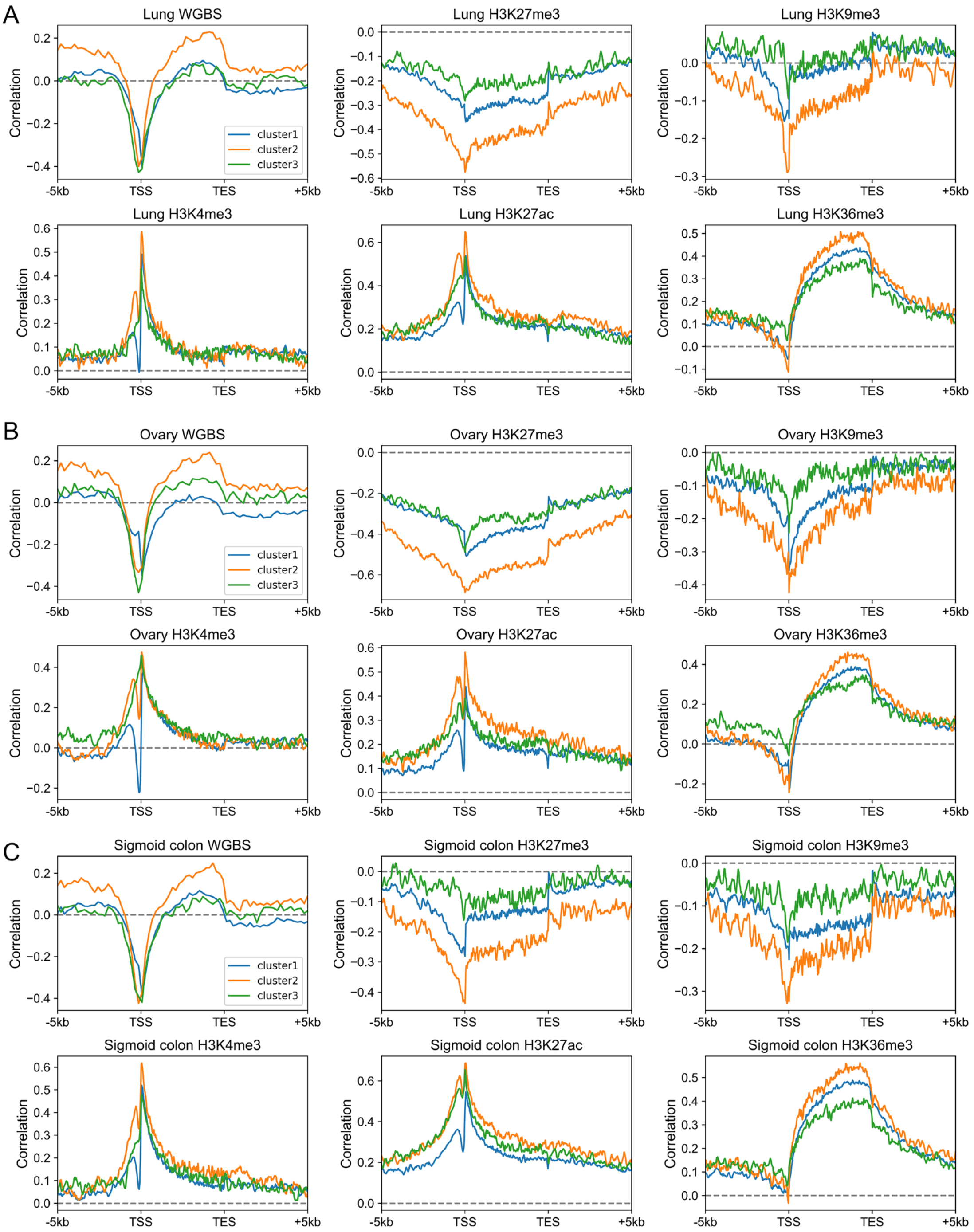


**Figure S7. The Spearman correlation between gene expression level and epigenetic marks in lung (A), ovary (B) and sigmoid colon (C).**

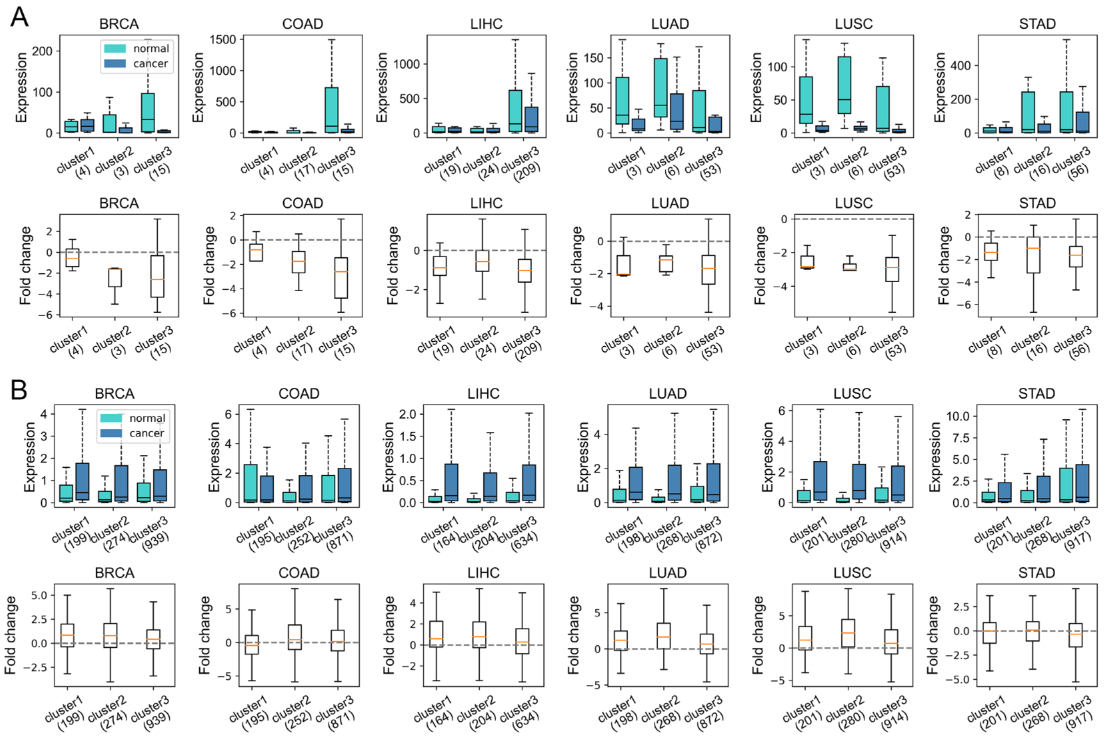


**Figure S8. (A)** The expression level (TPM) of tissue-specific genes (belonging to the corresponding normal sample) in normal and cancer samples, as well as the $\log_2$(expression fold change) in carcinogenesis calculated by Deseq2. **(B)** The expression level (TPM) of complementary tissue-specific genes in normal and cancer samples, as well as the $\log_2$(expression fold change) in carcinogenesis calculated by Deseq2.

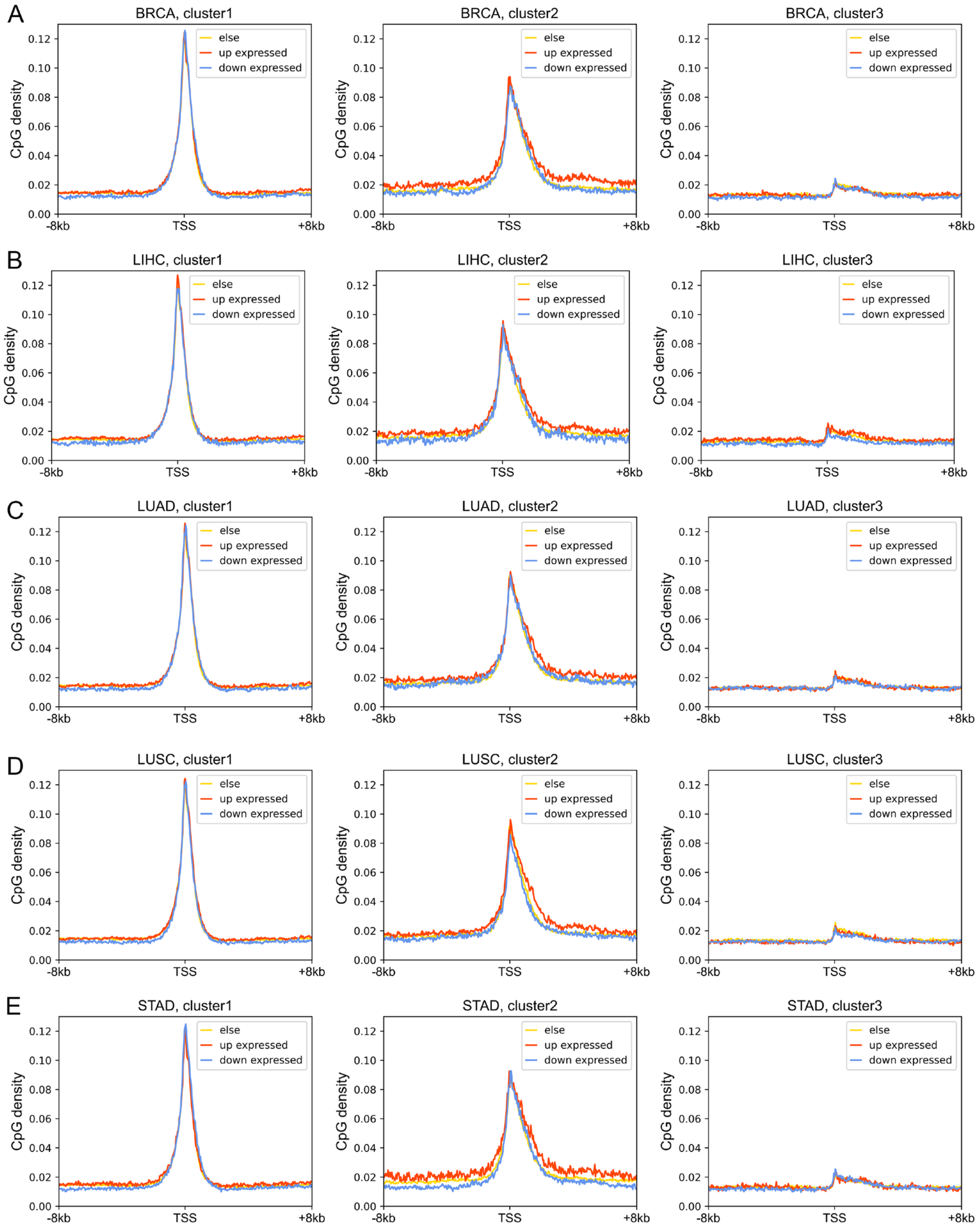


**Figure S9. The CpG density distribution of up-expressed, down-expressed and other genes.**

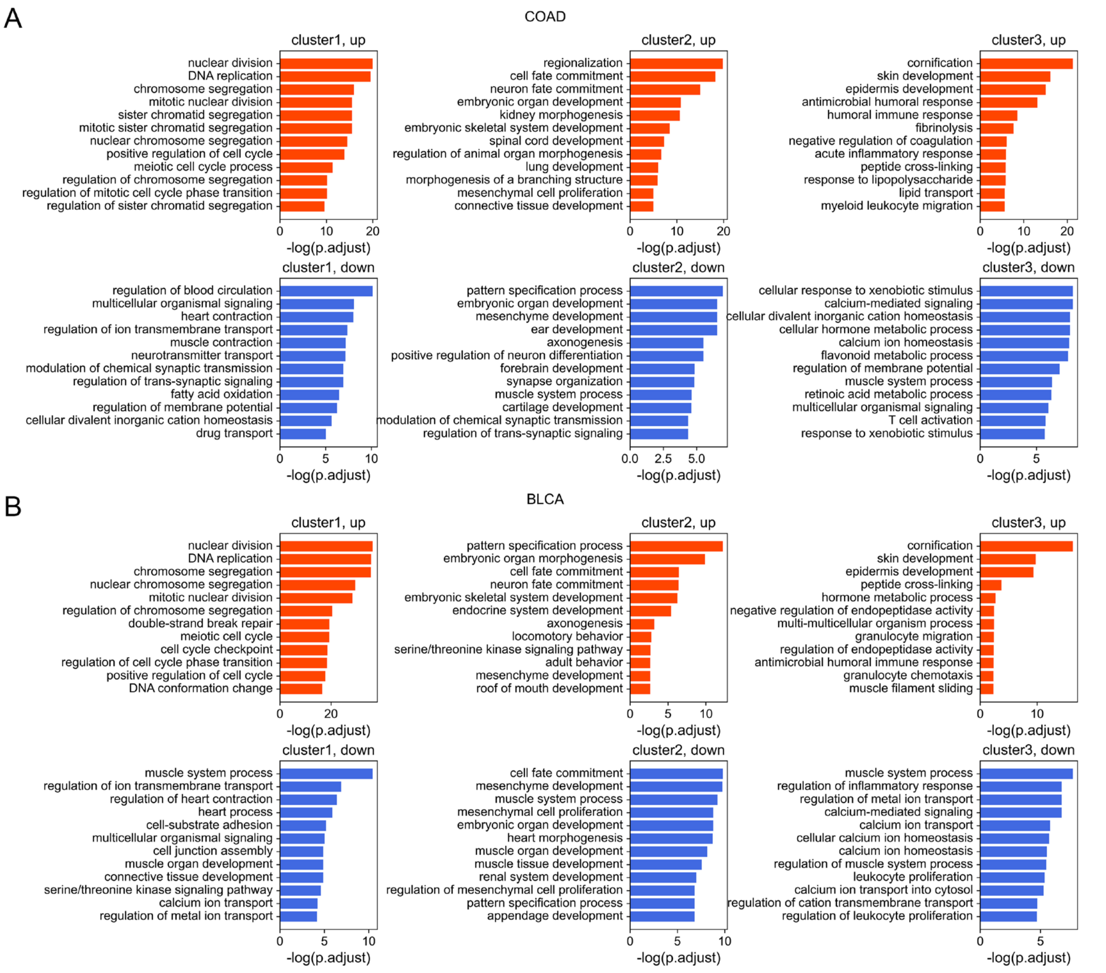


**Figure S10. The GO functions of up-expressed and down-expressed genes of different clusters.** COAD (colon cancer) and BLCA (bladder urothelial cancer) were used here for illustration.

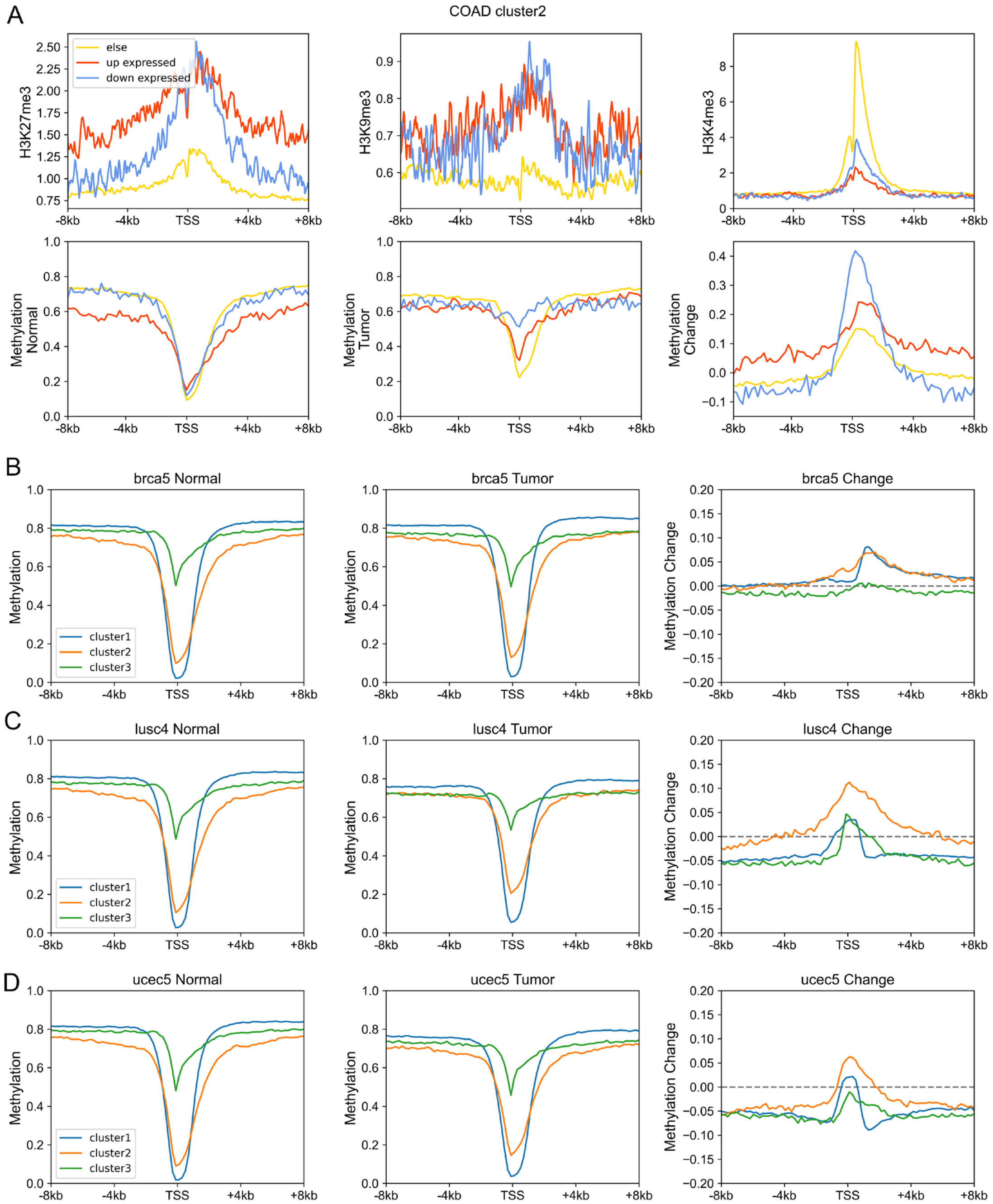


**Figure S11. (A)** Upper: the H3K27me3, H3K9me3, H3K4me3 of up-expressed, down-expressed and other genes within cluster 2 in corresponding normal cells. Down: the DNA methylation level of up-expressed, down-expression and other genes within cluster 2 in normal and tumor cells as well as the methylation change during carcinogenesis. **(B)-(D)** The DNA methylation level of genes of different clusters in normal (left) and tumor (middle) cells, and the methylation changes during carcinogenesis for three gene clusters (right).

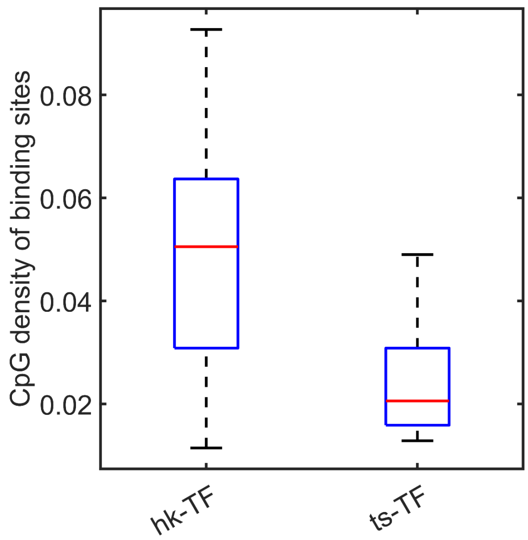


**Figure S12. The CpG density of housekeeping-TF binding sites and tissue-specific TF binding sites.**
